# Human Cervical Keratinocyte-Derived Monolayer and Organoid Cultures for Disease Modelling and Drug Screening

**DOI:** 10.1101/373456

**Authors:** Peter L. Villa, Robert Jackson, Statton Eade, Nicholas Escott, Ingeborg Zehbe

## Abstract

The successful isolation and propagation of patient-derived keratinocytes from cervical lesions constitute a more appropriate model of cervical disease than traditional cervical cancer-derived cell lines such as SiHa and CaSki. Our aim was to streamline the growth of patient-obtained, cervical keratinocytes into a reproducible process. We performed an observational case series study with 60 women referred to colposcopy for a diagnostic biopsy. Main outcome measures were how many samples could be passaged at least once, and where enough cells could be established, to precisely define their proliferation profile over time. Altering cell culture conditions over those reported by other groups markedly improved outcomes. We were also successful in making freeze backs which could be resuscitated for additional experiments. For best results, biopsy-intrinsic factors such as size and tissue digestion appear to be major variables. This seems to be the first systematic report with a well characterized and defined sample size, detailed protocol, carefully assessed cell yield and performance, and to successfully grow multi-layered, organoid cultures from cervical keratinocytes. This research is particularly impactful for constituting a sample repository-on-demand for appropriate disease modelling and drug screening under the umbrella of personalized health.

## Introduction

Primary neonatal foreskin keratinocytes are the most commonly employed human keratinocytes in the laboratory. Foreskins are easy to acquire, both practically and ethically speaking, and a foreskin can yield many more cells over a biopsy. Ideally, when studying cervical cancer, a researcher would use primary cervical cells, which can be acquired by isolating and culturing normal cervical keratinocytes from a healthy cervix and introducing the human papillomavirus (HPV) experimentally, or by propagating cells derived from naturally HPV-infected cervical lesion biopsies. Unfortunately, there are numerous practical and ethical hurdles when establishing cultures of normal primary cervical keratinocytes. First and foremost, it is unethical to acquire apparently healthy cervical tissue, as this introduces unnecessary risks, especially as many patients are of reproductive age. Therefore, the ideal means of isolating healthy cervical tissue is from a hysterectomy, which completely removes the uterus and cervix. Acquiring cervical tissue this way has many practical challenges, however, and requires the direct collaboration of the obstetrician/gynaecologist performing the procedure and the acting pathologist, which ensures the patient’s standard-of-care without formalin-fixing the still-living tissue.

The most practical and ethical approach seems to be obtaining biopsy specimens from patient lesions, while the physician is already taking a sample for standard-of-care. Given that cervical cancer results from persistent HPV infections, this approach would seem ideal for modelling and studying progression of cervical lesions. There are, however, numerous methodological hurdles to establishing an in vitro culture of keratinocytes from biopsy specimens. These issues can include, but are not limited to, microbial and fibroblast contamination^1^, low cellular viability and/or yield, and limited culture lifespan^2^. Previous studies demonstrated low colony forming efficiency^3^, limited potential to passage cultures^4^, and low adherence of viable cells^5^. Even with established methods, cultures from upper respiratory tract HPV lesions could be established from less than one-third of biopsy specimens and possessed a limited lifespan in vitro^2^. Currently, there is no literature on generating three-dimensional organotypic raft cultures (organoids) from patient-derived, naturally-infected cervical keratinocytes. Organoids have additional uses as they maintain many pathogen-host interactions while bypassing the need for animal models. Indeed, organoids have been used to test antiviral drugs against herpes viruses^6^ and poxviruses^7^, as well as topical treatments for head and neck squamous cell carcinoma^8^. It would seem feasible then to test new antiviral agents against HPV such as the ones developed by us^9^ in organoid models.

In this observational case series study, we developed a patient-based cell culture model using freshly excised, cervical biopsies as a more accurate alternative to traditional cervical cancer-derived cell lines such as SiHa and CaSki. Success was defined on how many samples could be propagated at least once, and then grown long enough to precisely define their proliferation profile over time and whether freeze backs could be made and resuscitated for additional experiments. Initial results based on published methods^1–5^ were clearly improved by altering cell culture conditions, and biopsy-intrinsic factors such as size and tissue digestion appear to be major variables. This seems to be the first systematic report with a clearly defined sample size, detailed protocol, carefully assessed cell yield and performance, and the ability to grow cervical organoids. To develop such an approach seemed meaningful for constituting a sample repository-on-demand for appropriate disease modelling and drug studies.

## Materials and Methods

### Study Design

We conducted a two-phase, observational case series study to develop a reproducible approach for growing cell cultures from human surgical specimen of the cervix uteri. The study was conducted at the Colposcopy Clinic located in the Thunder Bay Regional Health Sciences Centre and the associated Thunder Bay Regional Health Research Institute between August 2016 to March 2018. Fresh biopsies acquired from women attending the clinic were first taken for standard-of-care, followed by a swab for HPV typing and thereafter a secondary biopsy for the described research project (upon consent with one of the nurses on duty). We obtained altogether 60 biopsies: 30 for phase 1 and 30 for phase 2. The first 10 samples of phase 1 served to test the infrastructure and to compare the Stanley^1^ versus the Liu et al.^4,5^ method with 5 samples for each approach. As the Liu method was more successful in our hands, we continued employing it for systematic observations of the remaining 20 samples (Table 1). For phase 2, only 22 (all handled by the same investigator, P.L.V.) were used for the final evaluation to exclude experimental bias. The other 8 were handled by an investigator in training who since has left the institution.

**Table 1.** Successful establishment of keratinocyte cultures from cervical biopsies of phase 1. Cases follow in chronological order reflecting when biopsies were taken. Note that of 30 specimens acquired, only 20 are included in this table as explained in Materials and Methods. Only 3/11 specimens that initially yielded colonies were proliferating. Samples were divided into 4 groups: 0=No colonies of keratinocytes on starting in a T12.5 flask; 1=Visible colonies of keratinocytes but either no or too little growth to move to next size flask (i.e. from T12.5 to T25); 2=transfer to next size flask possible (T12.5 to T25); 3=transfer to next size flask possible (T25 to T75). Flask scaling: We typically split the T12.5 flask to the next size T25 flask once it was ~ 50% confluent because the centre of the colonies was typically quite dense; the T25 flask was trypsinized at ~ 80% confluence and all recovered cells were seeded on the subsequent T75 flask.

| Case ID | Category | Days to passage 1 | Days of life span | Size of biopsy |
| --- | --- | --- | --- | --- |
| 81 | 0 | - | - | Medium |
| 73 | 1 | - | - | Medium |
| 95 | 0 | - | - | Medium |
| 57 | 1 | - | - | Medium |
| 28 | 3 | 38 | 60 | Medium |
| 03 | 0 | - | - | Small |
| 68 | 0 | - | - | Large |
| 96 | 3 | 31 | 53 | Large |
| 39 | 1 | 18 | 30 | Small |
| 48 | 1 | - | - | Small |
| 17 | 1 | - | - | Large |
| 22 | 2 | 28 | 32 | Medium |
| 97 | 1 | - | - | Medium |
| 27 | 0 | - | - | Small |
| 43 | 1 | - | - | Medium |
| 59 | 1 | - | - | Large |
| 62 | 0 | - | - | Large |
| 84 | 0 | - | - | Small |
| 04 | 0 | - | - | Medium |
| 15 | 0 | - | - | Medium |

The only eligibility criterion for participation were women referred to the local Colposcopy Clinic and in need of a biopsy taken for diagnostic purposes because of previous abnormal Pap tests. Secondary punch biopsies of the endo- or ecto-cervix with areas suspicious of cervical neoplasia identified through colposcopy were used for this study. These were collected with Tischler Morgan biopsy forceps (3 mm x 7 mm bite) by the same colposcopist (N.E.) during the first scheduled visit after the primary biopsy was removed for diagnostic purposes. Biopsies were photographed with a ruler below them to measure their size.

### Outcomes and Parameters

We calculated how many samples could be passaged at least once, how many days this took, the overall life span (up to 60 days), and where applicable, the number of population doublings (PDs) and PD time (PDT) in hours. Cultures were grown until at least half of the cells had acquired a differentiated phenotype (enlarged cytoplasm with “fried egg” appearance). We also aimed to make freeze backs which could be resuscitated for additional experiments such as 3D culture. Biopsies varied between 1—4 mm^2^ and were classified as small, medium, or large. For phase 2 only, lesion grade, HPV status^10^ and patient age were obtained from the colposcopist (N.E.), while the researchers defined whether sample digestion prior to plating cells was good or poor.

### Monolayer (2D) Cell Culture

Initially, we compared the Stanley^1^ and Liu et al.^4,5^ method with 5 + 5 biopsies, respectively, to conclude which one to use for the remainder of the phase 1 samples. The “explant” method described by Stanley^1^ was discontinued because it yielded only sparse fibroblast colonies and we rather preferred to digest all the cells out. For the remaining 20 biopsies, we therefore switched to a modified version of the Liu et al.^4,5^ method: culture flasks were first coated with collagen (at least 30 min) with the Coating Matrix Kit (Fisher Scientific) and subsequently with fetal bovine serum (FBS) for 2—3 hours (rather than just coating in serum) to create a chemotactic surface. Biopsy specimens were placed in 5 mL of transport medium, consisting of serum-free Dulbecco’s modified Eagle’s medium (DMEM; Fisher Scientific) containing 20X antibiotic/antimycotic (Fisher Scientific) with 50 μg/mL gentamicin (Sigma Aldrich) and stored at 4°C for 1—4 hours until processing. Samples were then washed 5 times with 1 mL of Dulbecco’s phosphate-buffered saline (DPBS; Fisher Scientific) containing 20X antibiotic/antimycotic and 50 μg/mL gentamicin. Washed specimens were placed in 1 mL of 0.2% type I collagenase (Fisher Scientific) in 1X Hank’s balanced salt solution (HBSS; Sigma Aldrich) containing 3 mM calcium chloride and 1 mM magnesium chloride, without carbonate. Samples were minced with curved iris scissors and incubated on a shaker at 37°C and 225 rpm for 1 hour. Digested specimens were washed twice with 10 mL of DPBS, pelleted at 750 rpm for 5 minutes, and re-suspended in 2 mL of keratinocyte medium containing 5% fetal bovine serum (FBS; Fisher Scientific) and 1X antibiotic/antimycotic. Re-suspended tissue digests were plated onto a collagen- and FBS-coated T-12.5 flask as described above. After 4—6 days, the non-adherent tissue pieces were removed, and the medium was replaced with serum-free medium.

During phase 1, cervical cells were grown in keratinocyte serum-free growth medium (KGM; Cell Applications). In phase 2, we replaced the cell culture medium with EpiLife basal medium (Fisher Scientific) supplemented with human keratinocyte growth supplement (Fisher Scientific) as this formulation is rated to produce over twice as many population doublings^11^ thereby increasing proliferation potential. This is thought to be due to the decreased calcium concentration, the presence of insulin-growth factor (IGF)-I instead of insulin, and an increased number of organic components and inorganic salts. IGF-I possesses more mitogenic potential than insulin, as it activates more cellular receptors than insulin where IGF-receptors also interact with intracellular adaptor proteins more effectively than insulin receptors^12^. Moreover, IGF-I is more potent than insulin at activating MAPK in keratinocytes^13^ to stimulate proliferation via mitogenic signalling. To avoid stress signalling via the Rho pathway^14^, we also added the Rho kinase (ROCK) inhibitor Y-27632 (Fisher Scientific) to the medium. We reasoned this would further inhibit the differentiation of epithelial cells potentially resulting in increased keratinocyte proliferation^15,16^. Inhibition of ROCK reduces expression of down-stream targets for p53, which is thought to be a major mechanism by which Y-27632 promotes growth.

The immortalized mouse embryonic fibroblast cell line J2 3T3^17^ and primary human foreskin fibroblasts (ATCC, Cat. No. CRL-2097) were cultured in DMEM containing 10% heat-inactivated FBS and 1X antibiotic/antimycotic (Fisher Scientific). When passaging cultures, cell monolayers were washed with DPBS before adding 2 mL of 0.05% trypsin containing 0.02% ethylenediaminetetraacetic acid (trypsin-EDTA; Fisher Scientific) and incubated at 37°C for 5—10 minutes until cells were sufficiently detached, at which point 4 mL of culture medium was added to inactivate the trypsin. Cultures were seeded at 10% confluence and maintained up to 80% confluence before subsequent passaging. PDT was calculated by subtracting the difference between the log of the later population size from the log of the initial population size. The difference between the two values was divided by 0.301 (log of 2) to obtain the number of total PDs. PD time (hours) was then calculated by time elapsed divided by the number of PDs.

Keratinocyte cultures were maintained in a humidified incubator at 37°C with 5% CO2 at 10% to 80% confluence. Where applicable, cultures were split using 2 mL of trypsin-EDTA, whereby trypsin was inhibited with 3 mL of trypsin neutralizing solution (Cell Applications). The cells were centrifuged at 750 rpm for 5 minutes, where the neutralized trypsin solution was removed, and cells were re-suspended in fresh cell culture medium (i.e. KGM or EpiLife with Y-27632, phase 1 and 2, respectively). Cells were counted with a BioRad TC-10TM Automated Cell Counter. Keratinocyte cultures were regularly tested for mycoplasma indirectly using the J2 3T3 fibroblast cell line as an indicator cell line and DAPI nuclear stain^18^. Cells in culture were visualized using a Zeiss Axiovert 200 Inverted Microscope with an HBO 50 Watt AC-L2 mercury bulb connected to an MBQ 52 AC power supply. Images were taken with a QICAM 12-bit FAST 1394 camera. Phase-contrast was imaged at 100X magnification.

### Three-dimensional (3D) Organotypic Raft Cultures (Organoids)

Initially, the cervical rafts were grown with an in-house made medium as previously described for skin 3D organotypic rafts^19,20^. As this approach yielded less than expected epithelial multi-layering in the current study, we switched to a recently launched 3D medium specifically developed for growing rafts from gingival mucosal keratinocytes (CellnTec Advanced Cell Systems AG). This and doubling the number of fibroblasts showed the desired improvement in the growth process (multi-layered epithelium). Rafts were prepared in 3 to 6 technical replicates for each patient culture.

On day 1, low passage (P<5) human fibroblasts were used in the preparation of dermal equivalents. For 10 rafts, 0.8 x 10^6^ fibroblasts were re-suspended in 0.8 mL FBS. Using a sterile 10 mL beaker with a magnetic stir bar, cooled in ice atop a magnetic stir plate, 3 mL of type I rat tail collagen (~ 4.00 mg/mL; Millipore) and 400 μL of 10X HBSS was added and thoroughly mixed. The solution was neutralized using 5 μL increments of 5 N NaOH, requiring ~ 10—15 μL in total. Once neutralized, the fibroblasts were immediately added and thoroughly mixed. Dermal equivalents (DEs) were prepared by pipetting 400 μL of the collagen-cell suspension into wells of a 48-well plate. DEs were then incubated for 20 minutes at 37°C to solidify. Afterwards, 500 μL of DMEM-FBS was added on top of each DE and incubated overnight.

On day 2, for each raft, 0.25 x 10^6^ patient keratinocytes were re-suspended in 50 μL of EpiLife medium and seeded on top of the DE (to form the basal layer of the epidermis). After 2 hours, 500 μL of EpiLife with ROCK inhibitor was added to each raft and incubated overnight.

On day 3, the medium was removed, and rafts lifted to the air-liquid interface. A micro spoon was used to transfer the rafts from the 48-well plate onto Millicell 30 mm, 0.4 μm inserts (Millipore) placed in 6-well plates, where 1.1 mL of 3D medium was added around each insert, at one insert per well. The 3D medium was replaced every 2 days, where residual medium that accumulated around the rafts on top of well-insert membranes was removed. After 14 days, rafts were fixed overnight in 10% buffered formalin (Fisher Scientific). The rafts were subsequently washed once and stored in 70% denatured ethanol (Fisher Scientific) until processed further for histological assessment. We also characterized their p16 and cervical (mucosal) cytokeratin (CK) status (CK14, 17 and 19) by immunofluorescence as reported earlier by us for skin^20^. Primary antisera were rabbit monoclonals CK14 (1:2000; Abcam, ab181595), CK17 (1:150; Abcam, ab109725) and CK19 (1:500; Abcam, ab52625) as well as a mouse monoclonal p16 (1:100; Santa Cruz Biotech, sc56330). Hematoxylin & eosin images were taken at 200X magnification using a Zeiss Axioskop 50 with a QImaging Retiga 1300 camera. Immunofluorescence images were taken at 200X magnification with a QICAM 12-bit FAST 1394 camera.

### Statistical Methods

All statistical analyses were performed with the open-source statistical programming language R (version 3.5.0)^21^, within the integrated development environment RStudio (version 1.1.453)^22^. The significance level, alpha, was set to 0.05 for all analyses. Data were analysed using Fisher’s exact test for count data in phase 1 and multiple linear regression for phase 2.

### Ethics

This study was approved by the Research Ethics Boards (REBs) at Lakehead University (#019 16-17) and the Thunder Bay Regional Health Sciences Centre (TBRHSC; #2016107), according to Tri-Council Policy Statement 2 (2014) on human research ethics. Application to the REB required drafts of the: 1) logistical and laboratory protocols; 2) consent form; 3) brief study budget; 4) REB application template, involving risks and logistics; and 5) conflicts of interest. Phase I commenced in August of 2016 – over 8 months after registration.

For phase 2, which included patient chart review, altered laboratory protocols and HPV typing, an amendment was requested in June of 2017, after the study was renewed and the summary of the initial results was provided to the REBs. This amendment was granted through both institutions and the acquisition of samples for phase 2 began again in July of 2017.

## Results

From an original 60 cases (one biopsy per patient), 42 biopsies from women aged between 22 to 69 were finally evaluated for whether or not keratinocyte subcultures could be established, i.e. could be passaged at least once: a positive yield based on this criterion was obtained in 10/42 cases (~ 24%): 3/20 in phase 1 (15%) and 7/22 in phase 2 (32%). For phase 2, biopsies tended to be smaller in size, average days in culture until passage 1 was half that of phase 1 and longevity decreased by 25% (Tables 1, 2). All but 1 case were HPV-positive, 6 cases were lesion-free and approximately half digested well with collagenase (Table 2). Although reported earlier by another group^2^, none of the samples in either phase were contaminated throughout culturing.

**Table 2.** Summary of biopsy parameters, patient information, and culture growth of phase 2. Cases follow in chronological order reflecting when biopsies were taken. Biopsy size and visible digestion were noted on the day of sample processing. Patient’s age, HPV type(s), and lesion grade were recorded. Culture lifespan was measured as time of differentiation as in phase 1. Samples were divided into 5 groups: 0=No colonies of keratinocytes on starting in a T12.5 flask; 1=Visible colonies of keratinocytes but either no or too little growth to move to next size flask (i.e. from T12.5 to T25); 2=Transfer to next size flask possible (T12.5 to T25); 3=Transfer to next size flask possible (T25 to T75); 4=As 3 but at least 3 consecutive passages split at a ratio of 1:10 into a T75 to accurately assess PDTs. Flask scaling: We typically split the T12.5 flask to the next size T25 flask once it was ~ 50% confluent because the centre of the colonies was typically quite dense; the T25 flask was trypsinized at ~ 80% confluence and all recovered cells were seeded on the subsequent T75 flask.

| Case ID | Category | Days to passage 1 | Days of life span | Size of biopsy | Digestion | Lesion grade | HPV status |
| --- | --- | --- | --- | --- | --- | --- | --- |
| M33 | 0 | - | - | Medium | Poor | Metaplasia | 39 |
| UB1 | 0 | - | - | Small | Poor | CIN 1 | 45, 59 |
| 8N8 | 0 | - | - | Small | Poor | CIN 3 | 16 |
| QA7 | 0 | - | - | Small | Poor | Negative | 45 |
| A1U | 3 | 21 | 36 | Large | Good | CIN 1 | 6 |
| LY2 | 0 | - | - | Medium | Poor | CIN 1 | 53, 66 |
| HL7 | 0 | - | - | Medium | Poor | CIN 3 | 45 |
| IW9 | 3 | 19 | 24 | Small | Good | CIN 2-3 | 16 |
| 78I | 3 | 18 | 30 | Small | Poor | Negative | 52, 82, 89 |
| V7V | 0 | - | - | Small | Good | Negative | 16, 90 |
| 3UI | 4 | 13 | 50 | Medium | Good | CIN 2 | 31, 45, 89 |
| 04P | 2 | 20 | 28 | Medium | Good | CIN 1 | 51, 54, 81 |
| R4I | 0 | - | - | Small | Poor | Negative | 58 |
| AV6 | 4 | 9 | 26 | Medium | Good | CIN 3 | 16 |
| CS7 | 0 | - | - | Small | Good | CIN 1 | 33 |
| HW8 | 0 | - | - | Medium | Poor | CIN 1 | 45 |
| C5D | 1 | - | - | Large | Good | Negative | Negative |
| 7SG | 1 | - | - | Medium | Good | CIN 3 | 42, 52 |
| 83X | 1 | - | - | Medium | Poor | CIN 3 | 58, 90 |
| Z5X | 4 | 13 | 60 | Large | Good | CIN 2-3 | 16 |
| Y6A | 0 | - | - | Small | Good | CIN 2 | 18, 56, 81 |
| KR9 | 0 | - | - | Small | Good | CIN 1 | 6, 16, 70, 74 |

### Phase 1: Methods for Isolating and Propagating Cervical Keratinocytes

Biopsy specimens varied considerably in size. The smallest specimens (5/20) were approximately 1 mm in all dimensions and were too small to be minced. The largest specimens (5/20) were roughly a uniform 4 mm across. The remaining specimens (10/20) were in between these sizes (typically 2—3 mm wide and 1—2 mm in height and depth). The type of biopsy forceps used for excision depended on lesion size; hence, biopsy size varied according to lesion size and location, and the ease of accessing it with the forceps. There were notable differences in the ability to establish proliferative cultures from the various biopsy sizes (Table 1). The ability to establish growing cultures from biopsy specimens seemed to correlate with size as the 3 cases that could be passaged at least once either derived from medium or large biopsies. However, this finding was not statistically significant (Fisher’s exact test for count data, P = 0.850). Differentiation was evident upon microscopic examination, as cultures would exhibit noticeably reduced growth (i.e. modest changes in confluence over the course of one week). Cells would then cease to proliferate and markedly increase in cross-sectional area for approximately one or two more weeks (Figure 1). Therefore, lifespan was estimated retrospectively from the dates by which cells dramatically increased in size and had previously shown decreased proliferation (usually ~ 50% should have differentiated for the culture to be assessed as “differentiated”). Days in culture until passaged the first time was a mean of 32 days (range 28—38) and the average time until differentiation was 48 days (Table 1). An example of how keratinocytes grow out of digested tissue is displayed in Figure 2.

**Figure 1.**
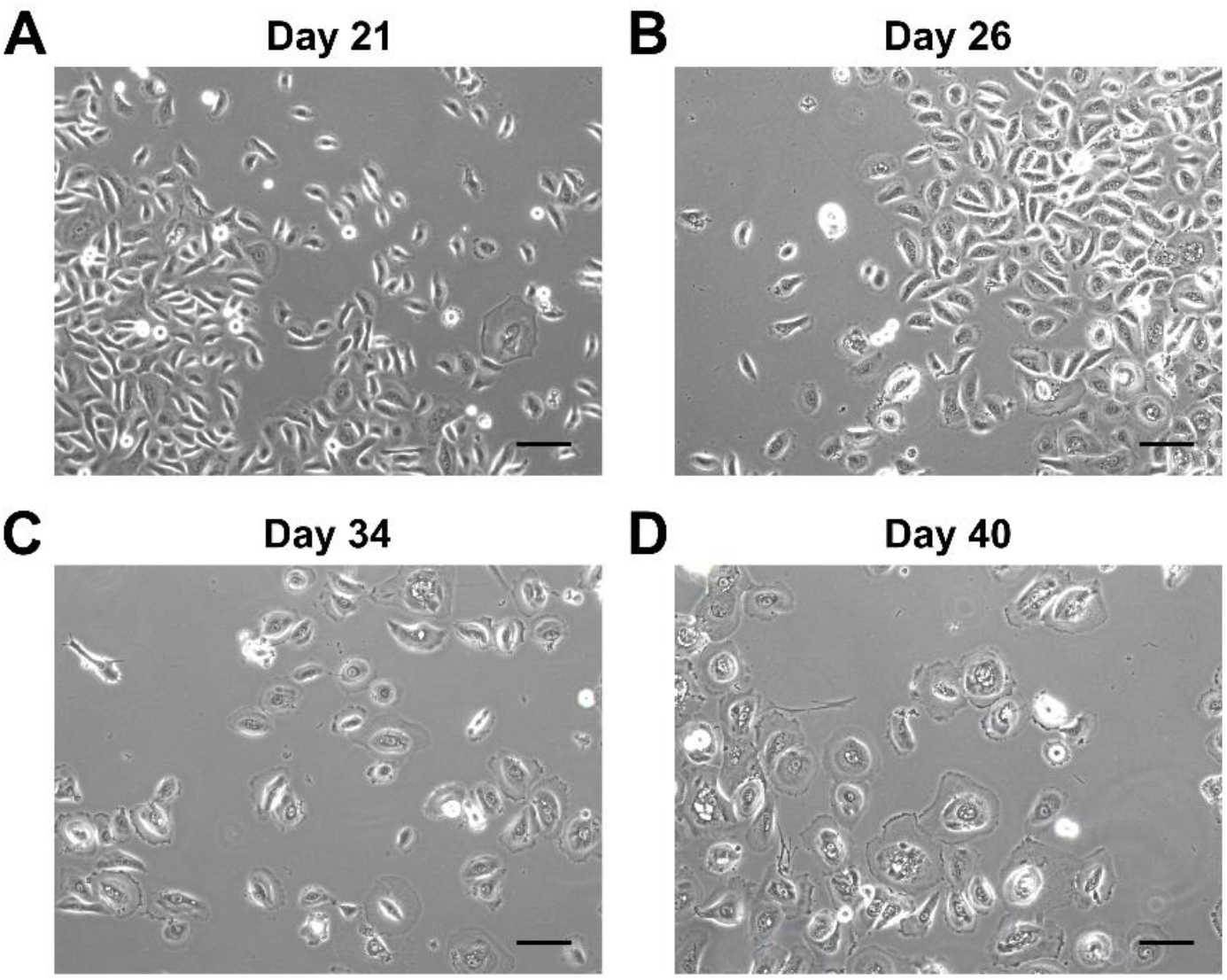
Visible and progressive differentiation of keratinocytes from patient isolate #22. In phase 1, numerical identifiers were used. Keratinocytes are visibly proliferating up to Day 21 (A). Keratinocytes have modestly but homogeneously increased in their apparent size by Day 26 (B). Keratinocytes are no longer proliferating, cell nuclei become centred, and cell membrane begins to spread along the flask evenly around the cell by Day 34 (C). Cells have uniformly increased in size by Day 40 (D). Magnification was 100X for A, B, C and D. Scale bars represent 100 μm.

**Figure 2.**
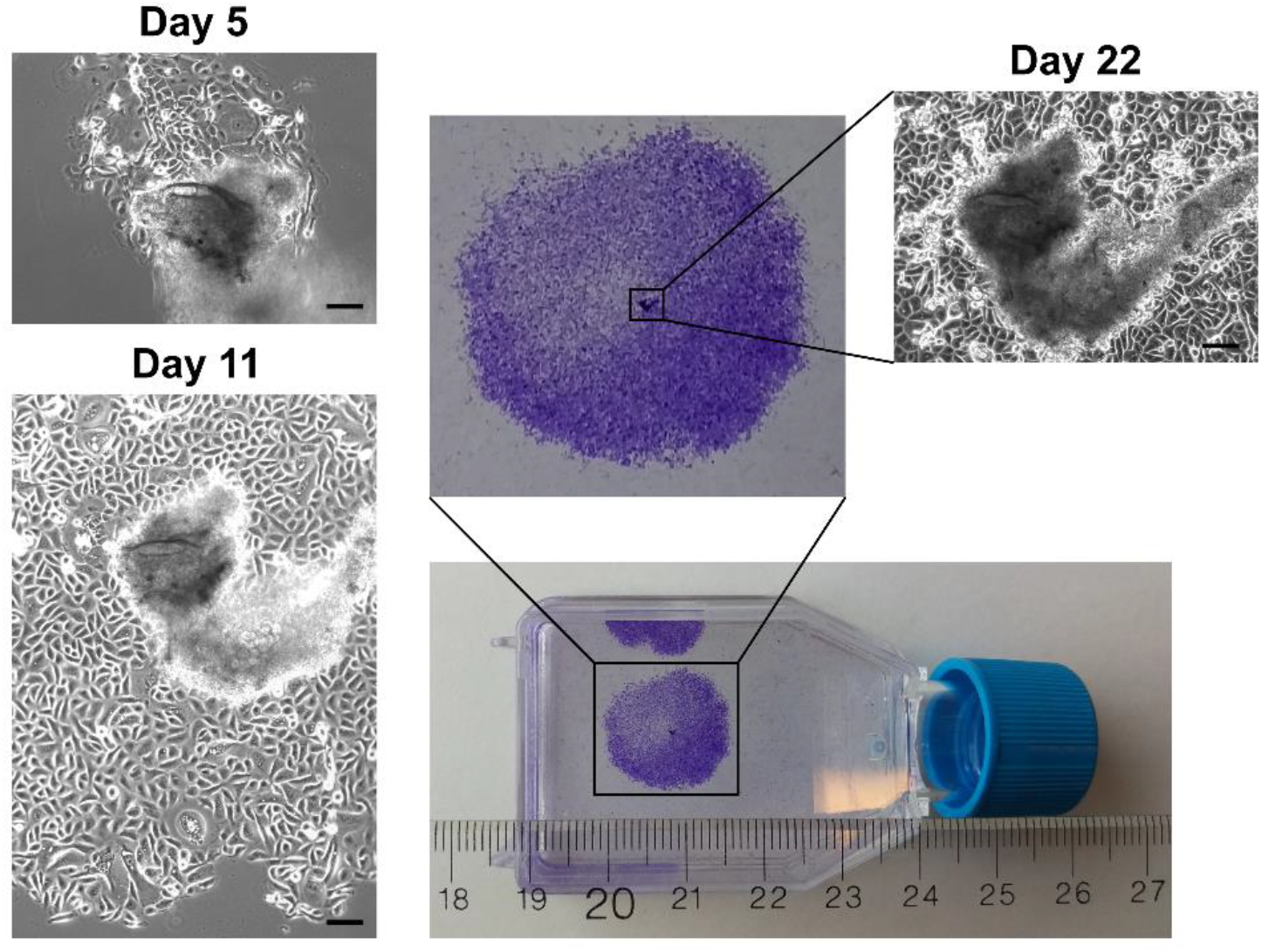
Time course of a single colony of cervical keratinocytes from patient isolate #59 (phase 1). Phase-contrast and crystal violet-stained monolayers of keratinocyte colonies growing out from fragments of partially digested tissue and behaving as pseudo-explant cultures. Days 5, 11 and 22 are detailed (100X magnification). Scale bars represent 100 μm.

### Phase 2: Significant Increase of Proliferation but not Longevity

Biopsy size metrics were the same as in phase 1. Size in phase 2 tended to be smaller at 9 small > 8 medium > 2 large versus 5 small > 10 medium > 5 large in phase 1 suggesting that our overall efficiency in establishing a proliferating culture was markedly improved by the new cell culture conditions (as explained in Materials and Methods). Additional patient information was collected. Patients had their cervices swabbed for HPV typing, and their age and lesion grade was recorded. From phase 1, it was also observed that some biopsy specimens would digest well with collagenase, where minced tissue fragments would digest into a visibly turbid cell suspension, and residual extracellular matrix would expand and clump, while others would not digest at all with collagenase. This observation appeared to be correlated with establishing colonies and therefore, in phase 2, it was noted whether each biopsy digested well or poorly as an additional parameter. Observations and results for establishing cultures are listed in Table 2. Days in culture until passaged the first time was a mean of 16 days (range 9—21) and the average culture lifespan was 36 days.

We assessed the predictive value of 4 independent variables: biopsy size, digestion, lesion grade, and HPV status. Most patients (82%) were found to be in their mid-twenties to mid-thirties, which we felt was too narrow a range as to include age as a separate variable. Multiple linear regression analysis revealed that a statistical model including biopsy size and digestion had a significant predictive value for biopsy success (P = 0.019, adjusted R^2^ = 0.359), whereas the inclusion of lesion grade and HPV status did not improve the model. There was a significant interaction effect between biopsy size and digestion (P = 0.041), where poor digestion was negatively associated with success in medium-sized biopsies (P = 0.041), and small biopsies had the lowest overall success (P = 0.037). There were 3 patient cultures that proliferated long enough to have their population doubling times (PDTs) accurately calculated via cell counts, test cryogenic storage and grow rafts; samples 3UI, AV6 and Z5X all could be passaged more than 3 times on T75 flasks yielding 3—5 million cells amounting to ~ 3 population doublings (PDs) per passage. Freeze backs were made from the first ~80% confluent T75 flask. Average population doubling time (PDT) was 27 hours (range 23—30): 3UI (30), AV6 (23) and Z5X (27). Passages and PDs were calculated when harvesting the first (subtracting the days from total lifespan) and subsequent ~ 80% confluent T75 flasks (until onset of differentiation). Average number of passages was 8 (range 5—11): 3UI (7), AV6 (5) and Z5X (11). Average total PDs until differentiation was 24 (range 15—32): 3UI (26), AV6 (15) and Z5X (32).

### Organotypic Raft Cultures

When seeding keratinocytes onto dermal equivalents, the size of the keratinocytes was noted from the TC-10TM Cell Counter. Typically, keratinocytes were homogeneously between 16 and 18 μm in diameter while actively proliferating. When cells were beginning to differentiate, a discrete proportion of the population would exist between 22—24 μm. For example, when 30—50% of the cells were 22—24 μm, the culture would differentiate on the current passage when re-seeded. This observation held true for all cultures in phase 2. The proportion of small-to-large cells was recorded upon passaging to note when cultures would differentiate. An attempt was also made to grow raft cultures from cells that could only be passaged once or twice onto T75 flasks, i.e. for A1U, IW9, and 78I. It was noted for these patient cultures that one-third of the population existed at 22—24 μm, illustrating that they were near the end of their in vitro lifespan at the time they were seeded onto dermal equivalents. Indeed, the remaining cell suspension (i.e. in excess from seeding for raft cultures) were re-seeded back into monolayer culture and differentiated without further passaging. Rafting did not succeed in these cases where only a couple of layers would form. In contrast, 3UI, AV6, and Z5X could all be grown as rafts (Figure 3) and showed homogeneously small cells (i.e. 16—18 μm) at the time of 3D culturing. Hematoxylin and eosin (H & E) staining displayed dysplastic epidermis with up to 8 keratinocyte layers. To determine epithelial differentiation, rafts were immunostained with cytokeratins (CKs) normally seen in cervical epithelium: CK14, 17 and 19 staining resembled an abnormal pattern typically found in high-grade cervical lesions^23^. As a proxy biomarker, diffuse cytoplasmic p16 staining was indicative of a high-risk HPV infection.

**Figure 3.**
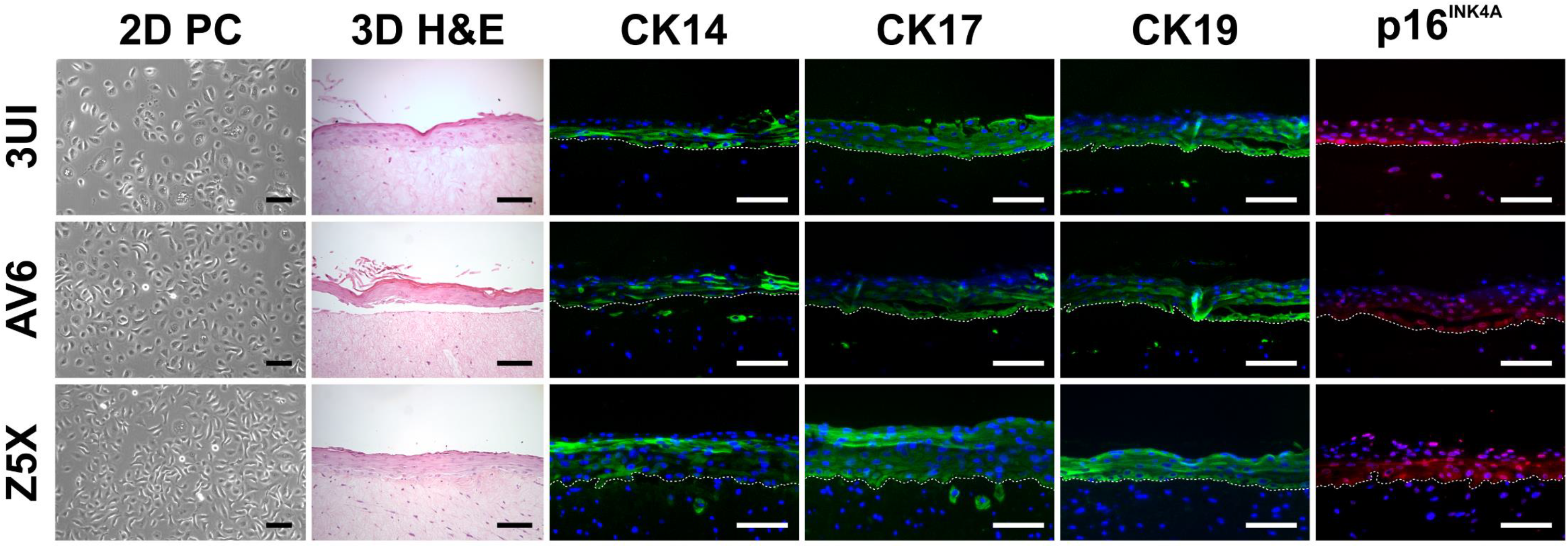
Successful biopsy-derived isolates 3UI, AV6 and Z5X (phase 2). From left to right: Phase-contrast micrographs of keratinocyte monolayers (100X magnification), hematoxylin and eosin stained micrographs of sectioned formalin-fixed paraffin-embedded raft cultures (200X magnification), and immunofluorescence micrographs of cytokeratin (CK) 14, 17, 19, and p16^INK4A^ of sectioned formalin-fixed paraffin-embedded raft cultures (200X magnification). Green fluorescence represents cytokeratins; red fluorescence represents p16^INK4A^; blue DAPI staining indicates nuclei; white dashed line represents basal cell membrane intersecting epidermis and dermis (from top to bottom). Scale bars represent 100 μm.

## Discussion

With the current investigation, we have provided a detailed report of culturing cervical keratinocytes from patient biopsies rather than hysterectomies. We could successfully expand 3 individual cultures and grow organotypic rafts after resuscitation. Our current protocol should generate enough keratinocytes for an appropriate amount of independent experiments with multiple replicates from various donors to allow robust biomarker, statistical and next generation sequencing analyses^19^. Successful culturing largely depended on the nature of the biopsy with regard to its size and digestion ability before plating. An interesting insight regarding the latter was recently published by Fan and colleagues^24^ finding that enzymatic dissociation of keratinocytes worked better with dispase II combined with 0.25% trypsin–0.01% EDTA than with collagenase I. This may be due to the relative content of stromal components in cervical biopsies such as fibronectin versus collagen.

It is now possible to purchase hysterectomy-derived cervical keratinocytes from at least 3 independent vendors. An inquiry, with one of them replying, revealed that they guarantee 3 passages with a total of 10 PDs and a PDT typically between 40 to 70 hours (depending on medium composition). In the Fan and colleagues’ study^24^, where keratinocyte cultures were likewise obtained from hysterectomies, 5—6 passages with 1 PDT per passage was reported. Taken together, this suggests that our samples outperform both these accomplishments despite the fact that they originated from biopsies: an average of 8 passages, each with 3 PDs, and a PDT of 27 hours, expecting to yield several hundred million cells while also preparing cell stocks. Other studies did not report their data in a similarly quantitative manner^1–5^, precluding any meaningful comparison with our results. The current study seems to be stand-alone to systematically investigate the culturing of biopsy-derived, cervical keratinocytes both as monolayers and organotypic rafts. We also report success and shortcomings in a descriptive and quantitative, open-ended manner implying that there is still room for improvement.

The biggest variance between phase 1 and phase 2 was the proliferation efficiency, which is likely an effect from our adopted medium condition with IGF-1 and ROCK inhibitor components. While both components may have helped to increase proliferation of progenitor cells short term, they may cause “proliferative exhaustion” after an initial enhancement due to an increase in mitosis. Long-term, however, exposure to IGF-1 has been shown to induce premature senescence, e.g. through inhibition of SIRT1-mediated p53 deacetylation^25^, which may explain the somewhat shorter temporal lifespan in phase 2 compared to phase 1. Boosting stem cells that potentially reside in the obtained biopsies will therefore be considered as a next step to generally increase the yield of cultures. Indeed, stem cells may be propagated with specific medium formulations supplemented with factors such as Wnt, Noggin, R-Spondin and EGF and only the latter is present in any of the media we used in this study.

Experiments are underway to investigate whether an IGF-1 weaning off by switching back to classical KGM will increase temporal lifespan of cervical keratinocytes. The ROCK inhibitor’s short- and long-term effect will likewise be further investigated. We included it in this study to complement the effect of IGF-1, and used it without fibroblast feeders to boost proliferation without inducing immortalization^15^. We will also test whether low oxygen concentration helps further increase temporal lifespan^26^. Most keratinocyte culture experiments are conveniently performed using neonatal human skin keratinocytes. It remains to be fully determined how lessons learned from such studies apply to adult mucosal keratinocytes.

Acquiring keratinocytes through cervical biopsies rather than hysterectomies may be a tedious and time-consuming endeavour yet is essential to capture a close-to-diagnosis scenario. Our group wanted to create a better than the traditional and exhausted carcinoma-derived cell lines model for our in-house therapy development^9,27^. With these prospects in mind, we designed the present study, but alternatives should also be considered. There is new evidence that patient-derived material has its own drawbacks. In a recent Nature Genetics report, the ever-changing evolution of patient-derived material due to artificial cell culture-specific conditions was discussed^28^. Based on 3 types of cancer models—patient-derived xenografts, carcinoma-derived cell lines and xenograft-derived cell lines, the authors concluded that switching the environment in which a model is propagated results in “tumour dynamics” that gradually alter its genetic landscape. All cancer models are potentially affected by clonal selection suggesting that genomic instability and tumour heterogeneity have been underappreciated, impacting the approach to target individual biomarkers in the personalized health landscape.

It needs to be established if mucosal, patient-derived keratinocytes from pre-malignant lesions such as in our scenario are equally prone to the above-mentioned tumour dynamics as fully malignant and invasive tumours are. This would require getting multiple biopsies from patients diagnosed with cervical dysplasia, which may pose an ethical issue, and compromise patient safety as well as diagnostic and treatment regimens of standard-of-care.

It also needs to be investigated if the targeting of high-risk HPV expression would lead to a beneficial change in phenotype from that of a proliferating tumour to that of tumour elimination by apoptosis as HPV seems to be required for the malignant phenotype at least in early tumourigenesis as generally hypothesized^9^. The presence of HPV genomic DNA and its reported variability both impact on the malignant transformation of keratinocytes, the host cell of HPV. Genomic instability is a frequent phenomenon in HPV-positive keratinocytes and differs in various geographic sub-lineages as reported by us^19,20^. Next steps by our lab are therefore to target HPV rather than individual host biomarkers by eliminating HPV oncogene expression on the mRNA^9^ and protein level^27^. Aside from our goal to launch a cell repository for future pre-clinical trials based on patient biopsies, we will also use commercially obtained, cervical keratinocytes using our here reported, more efficient cell culture protocol. Such an engineered model with varying amounts of HPV genome copies based on the content in real human dysplastic lesions may be useful for more experimental-oriented work^19,20^. This HPV-focused approach will not require lengthy ethical processes and can be adapted to any in vivo HPV context.

## Author Contributions

For this research article, individual contributions were as follows in order of magnitude: Conceptualization, I.Z., P.L.V., R.J. and N.E.; Methodology, P.L.V., I.Z., N.E.; Software, R.J.; Validation, R.J., I.Z., P.L.V., S.E., N.E.; Formal Analysis, R.J., I.Z., P.L.V.; Investigation, P.L.V., N.E., I.Z.; Wet lab work: P.L.V., I.Z., S.E.; Resources, I.Z.; N.E.; Data Curation, P.L.V., N.E.; Writing – Original Draft Preparation, I.Z., P.L.V.; Writing – Review & Editing, I.Z., R.J., P.L.V., S.E., N.E.; Visualization, S.E., R.J., I.Z.; Supervision, I.Z., N.E.; Project Administration, P.L.V., I.Z.; Funding Acquisition, I.Z.

## Funding

This research was funded by a Collaborative Health Research Projects (CHRP) grant between the Canadian Institutes of Health Research (CIHR) and the Natural Sciences and Engineering Research Council of Canada (NSERC) to I.Z. (CIHR: CPG 140188 and NSERC: CHRP 478520-15); NSERC graduate student scholarships to P.L.V. and R.J. (CGS-D3: 454402-2014) as well as a Northern Ontario Heritage Fund Corporation (NOHFC)-sponsored internship for S.E.

## Acknowledgments

The authors acknowledge the excellent assistance of the study nurses Ms. Bonnie Kozak, Ms. Marita Mauro and Ms. Brenda Kulifaj in enrolling patients as well as of Dr. Alberto Severini in performing HPV testing and typing.

## Conflicts of Interest

The authors declare no conflict of interest. The founding sponsors had no role in the design of the study; in the collection, analyses, or interpretation of data; in the writing of the manuscript, and in the decision to publish the results.

